# Ubiquilin proteins regulate EGFR levels and activity in lung adenocarcinoma cells

**DOI:** 10.1101/2020.06.03.131888

**Authors:** Zimple Kurlawala, Kumar Saurabh, Rain Dunaway, Parag P. Shah, Leah J. Siskind, Levi J. Beverly

**Affiliations:** James Graham Brown Cancer Center, School of Medicine, University of Louisville, Louisville, KY USA; School of Medicine, University of Louisville, KY, USA; Department of Pharmacology and Toxicology, University of Louisville, Louisville, KY, USA; Division of Hematology and Oncology, School of Medicine, University of Louisville, Louisville, KY, USA; Laboratory of Molecular Gerontology, National Institute on Aging, National Institutes of Health, Baltimore, MD, USA

**Keywords:** UBQLN1, Ubiquilin, EGFR, cancer, IGFR

## Abstract

Ubiquilin proteins (UBQLNs) are involved in diverse cellular processes like ERAD (endoplasmic reticulum associated degradation), autophagy, apoptosis and epithelial to mesenchymal transition. UBQLNs interact with a variety of substrates, including cell surface receptors, transcription factor regulators, proteasomal machinery proteins, and transmembrane proteins. Additionally, previous work from our lab shows that UBQLN1 interacts with IGFR family members (IGF1R, IGF2R, INSR) and this interaction regulates the activity and proteostasis of IGFR family members. Here, we examined regulation of UBQLN1 with Epidermal Growth Factor Receptor (EGFR) in lung adenocarcinoma cells. Loss of UBQLN1 occurs at high frequency in human lung cancer patient samples and we have shown that loss of UBQLN1 is capable altering processes involved in cell proliferation, migration, invasion and epithelial to mesenchymal transition in lung adenocarcinoma cell lines. Here, we present data that loss of UBQLN1 resulted in increased turnover of total EGFR, whilst increasing the relative amount of active EGFR in lung adenocarcinoma cells, especially in the presence of its ligand EGF. Furthermore, loss of UBQLN1 led to a more invasive cell phenotype as manifested by increased proliferation, migration and speed of movement of these lung adenocarcinoma cells. Taken together, UBQLN1 regulates expression and stability of IGFRs and EGFR, members of the receptor tyrosine kinase family of proteins in lung cancer cells.

## Introduction

Cancer and Alzheimer’s disease (AD) are seemingly caused by contrasting cellular processes; aberrant cell survival for cancer and aberrant cell death for AD (Shafi, 2016). The family of adapter proteins, Ubiquilins (UBQLNs), are lost in multiple types of cancers as well as in AD (Beverly, Lockwood, Shah, Erdjument-Bromage, & Varmus, 2012; Viswanathan et al., 2011; Y. Wang et al., 2015). This family consists of five members; UBQLN1-4 and UBQLNL, which contains an N-terminus ubiquitin-like (UBL) domain and a C-terminus ubiquitin-associated (UBA) domain (Kleijnen et al., 2000; Z. Kurlawala, Shah, Shah, & Beverly, 2017; Marin, 2014). Ubiquilin1 is involved in a variety of cellular processes like ERAD (endoplasmic reticulum associated degradation) (Lim et al., 2009; Shah et al., 2015), autophagy (Lee, Arnott, & Brown, 2013; Elsa-Noah N’Diaye et al., 2009), apoptosis (Sun et al., 2015) and epithelial to mesenchymal transition (EMT) (Shah et al., 2015; Yadav et al., 2017). Ubiquilin1 also interacts with diverse substrates – proteins involved in the proteasomal machinery (PSMD4, BAG6) (Z. Kurlawala, Shah, et al., 2017) cell surface receptors, GABA-A (Saliba, Pangalos, & Moss, 2008), GPCR’s (E. N. N’Diaye et al., 2008), PSEN1/2 (Mah, Perry, Smith, & Monteiro, 2000; Massey et al., 2004), IGF1R (Z. Kurlawala, Dunaway, et al., 2017; Z. Kurlawala, Shah, et al., 2017), transcription factor regulators, IκBα (Feng et al., 2004) and other transmembrane proteins ESYT2 (Z. Kurlawala, Shah, et al., 2017), CD47 (Wu, Wang, Zheleznyak, & Brown, 1999) and BCLb (Beverly et al., 2012). Ubiquilin1 is a versatile, multi-purpose adaptor that interacts with a wide range of substrates; thus, it can regulate multiple important cellular processes.

UBQLN2, is another UBQLN family member that is constitutively expressed in most cell types and shares more than 75% homology with UBQLN1, indicating that it likely shares similar biological functions(Marin, 2014). Like UBQLN1, UBQLN2 also has a UBL domain which interacts with the proteasome and a UBA domain which recognizes ubiquitin on target proteins (Kleijnen, Alarcón, & Howley, 2003; Renaud, Picher-Martel, Codron, & Julien, 2019). Additionally, UBQLN2 has a 12-PXX repeat region which makes it unique among the UBQLN family proteins (Renaud et al., 2019).

Receptor tyrosine kinases (RTKs) are cell surface receptors found to be responsible for mediating signaling pathways crucial to cell proliferation, cell migration and invasion of many types of cancer (Zwick, Bange, & Ullrich, 2001). Our lab was first to identify interaction of UBQLN1 with a RTK family member, namely insulin-like growth factor receptors (IGFRs) (Z. Kurlawala, Dunaway, et al., 2017). Loss of UBQLN1 leads to a significant decrease in the amount of total IGF1R, an increase in phosphorylated IGF1R and dramatic increases in their migratory potential when stimulated with IGF in lung adenocarcinoma. Epidermal growth factor receptor (EGFR), a transmembrane protein that is also a member of the RTK family, is mutated or over-expressed in multiple cancers and AD (Lurje & Lenz, 2009; Porta et al., 2011; Tavassoly, Sato, & Tavassoly, 2020). EGFR is one of the most commonly studied oncogenes to date. It is often upregulated in multiple cancers, and downregulated in AD (Shafi, 2016). This unique inverse relationship between cancer and AD has been classified for a wide array of proteins, including p53, IGF1R and BCL2 (Shafi, 2016). EGFR is a well-established therapeutic target of kinase inhibitors (gefitinib, erlotinib, and lapatinib) and monoclonal antibodies (cetuximab, panitumumab, and trastuzumab). Many of these therapeutics are now being used as first line treatment for both lung cancer and Alzheimer’s patients (Lurje & Lenz, 2009; Porta et al., 2011; Tavassoly et al., 2020). In this study, we present data establishing the interaction of UBQLN1 and UBQLN2 with EGFR. In lung adenocarcinoma cell lines, downregulation of UBQLN1, followed by EGF stimulation, leads to degradation of total EGFR protein, and an increase in migration and invasion potential of these cells.

## Materials and Methods

### Cell Culture, Transfection and EGF stimulation

Human embryonic kidney 293T (HEK293T) cells were procured from American Type Culture Collection (ATCC, Rockville, MD, USA) and cultured in DMEM medium (#SH30243, Hyclone, Logan, UT, USA) supplemented with 10% fetal bovine serum (#SH30070, Hyclone, Logan, UT, USA) and 1% antibiotic/antimycotic (#SV30010, Hyclone, Logan, UT, USA) at 37°C with 5% CO2. A549 (lung adenocarcinoma line) were procured from ATCC and cultured in RPMI (#SH30027, Hyclone, Logan, UT, USA) supplemented with 10% FBS, 1% antibiotic/antimycotic. siRNA transfections were performed as described previously(Z. Kurlawala, Dunaway, et al., 2017).^19^ Briefly, 48 hours after transfection, cells were serum starved (SS) for 3 hours, incubated with protein synthesis inhibitor, Cycloheximide (CH, 10μM) for 1 hour prior to supplementing serum-free media with 25 or 50 ng/ml based on the experiment. EGF (#PHG0314, Thermo Fisher Scientific, Waltham, MA, USA). 3 hours later, cells were harvested and probed for EGFR (total and phosphorylated) protein by Western Blot analysis. For dose and time-dependent studies, cells were stimulated and/or incubated for doses and time-points indicated in respective figures.

### Plasmid Construction

As described previously, constructs with deleted domains of UBQLN1 (Figure 1A), were developed using Q5 Site-Directed mutagenesis kit as per manufacturers protocol (New England Biolabs # E0554; Ipswich, MA, USA) and confirmed by sequencing(Z. Kurlawala, Shah, et al., 2017).^7^

**Figure 1:**
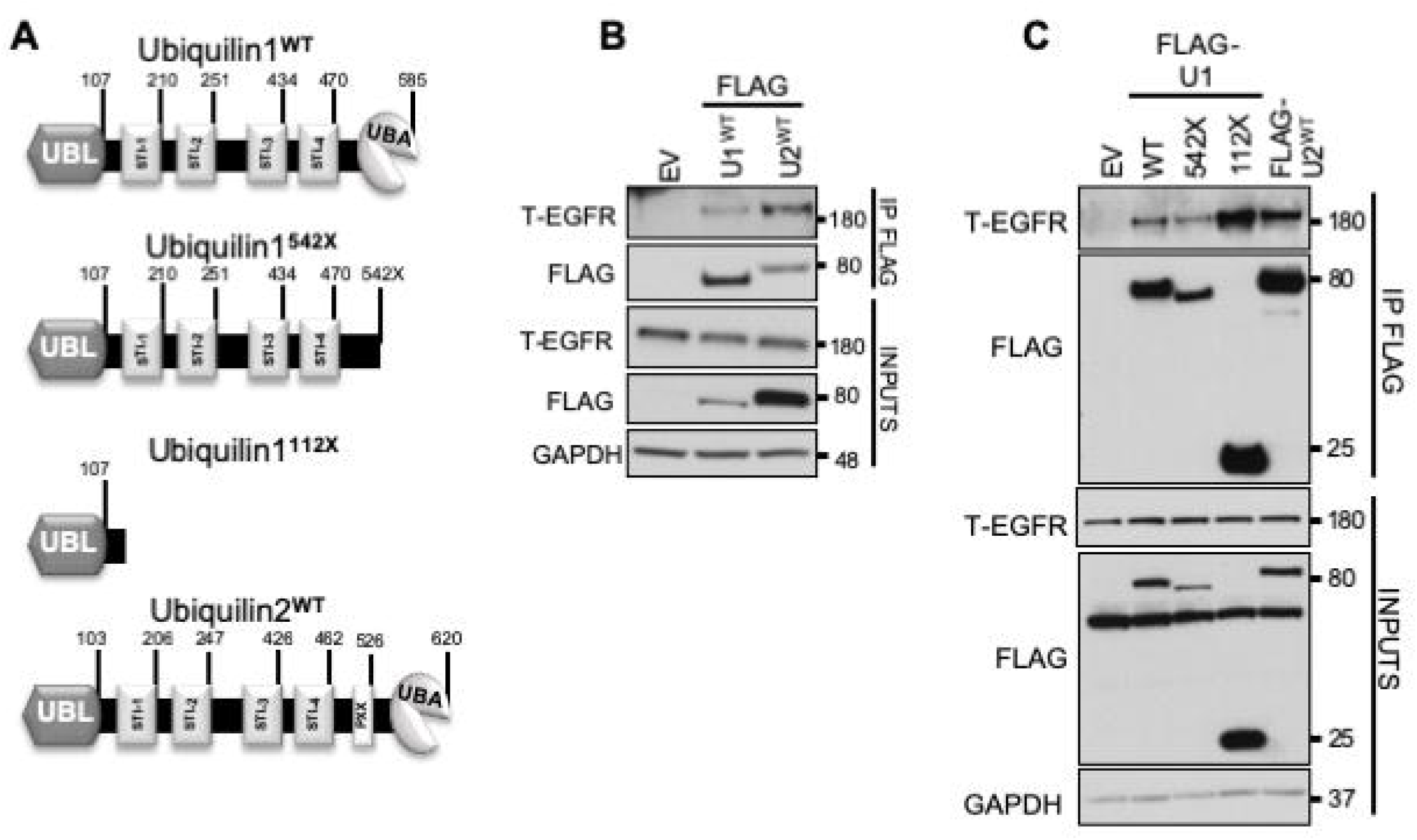
Ubiquilin1 and Ubiquilin2 interact with EGFR. **(A)** Schematic of Ubiquilin1^WT^, Ubiqulin1^542X^, Ubiquilin1^112X^ and Ubiquilin2^WT^ constructs. Ubiquilin1 (590 amino acids) and Ubiquilin2 (620 amino acids) proteins have an N-terminal UBL domain, four STI chaperone-like domains in the middle and a C-terminal UBA domain. Ubiquilin2 has an additional 12-PXX repeat region. **(B)** HEK293T cells were transiently transfected with FLAG-tagged Ubiquilin1 (U1) and Ubiquilin2 (U2) followed by co-immunoprecipitation (IP) by anti-FLAG antibody and Western Blot analysis for total EGFR. Both Ubiquilin1 and Ubiquilin2 interact with T-EGFR. **(C)** HEK293T cells were transiently transfected with FLAG-U1^WT^, FLAG-U1542X, FLAG-U1112X or FLAG-U2 followed by co-immunoprecipitation by anti-FLAG antibody and probed for EGFR. All 3 constructs of U1 interact with T-EGFR, indicating that the UBA domain is dispensable for interaction between these two proteins.

### Immunoprecipitation, Protein estimation and Western Blot

Immunoprecipitation (IP) was performed as described previously. Harvested cells for each procedure (IP and/or transfection) were lysed with 1% CHAPS lysis buffer and total protein was estimated using the BCA quantification method(Z. Kurlawala, Shah, et al., 2017). Western blot analyses were performed in Bolt Bis-Tris gels (#BG4120BOX, Life Technologies, Grand island, NY, USA) as per manufacturer’s protocol using antibodies from Santa Cruz, Dallas, TX, USA (GAPDH # sc47724); Sigma-Aldrich, St. Louis, MO, USA (Actin # A5316); Yenzym Antibodies LLC, South San Francisco, CA, USA (UBQLN polyclonal was made by inoculating rabbits with a peptide specific to UBQLN1); and Cell Signaling, Danvers, MA, USA (UBQLN1 # 14526, FLAG # 14793, EGFR # 4267, pEGFR Tyr1068 # 3777).

### Cell viability and Migration assay

Cell viability and migration assays were performed as described earlier.^19^ Briefly, A549 cells were cultured in 60mm culture plates. After 12 hours of transfection with siRNA, cells were trypsinized, counted and 2000 cells were reseeded per well in 96-well plates in complete media. 12 hours post-reseeding in complete media, cells were serum starved for 3 hours followed by stimulation with EGF and were cultured in media containing 2% FBS. Cell viability was analyzed for four successive days using AlamarBlue^™^ (#DAL1100, Thermo Fisher Scientific, Waltham, MA, USA). At the same time following transfection, 5000 cells were seeded in Transwell^™^ cell culture inserts (#CLS3464, Corning Inc., Corning, NY, USA) in triplicate for each condition as described previously.^19^ Briefly, cells were allowed to grow on Transwell^™^ cell culture insert in serum free media, serum free media supplemented with EGF (50ng/ml) and serum free media supplemented with both EGF and Erlotinib (1μM). After 24hrs, membranes were washed once with PBS, fixed with ethanol, stained with Giemsa stain (#R03055, Sigma-Aldrich, St. Louis, MO, USA) and cells were counted on microscope.

### Live Cell Imaging

A549 cells were transfected with siRNA against UBQLN1 and non-targeting control. 24 hours post-transfection, cells were trypsinized, counted and 10,000 cells were reseeded per well in 12-well plate, coated with thin layer commercial extracellular matrix (ECM # E6909, Sigma-Aldrich, St. Louis, MO, USA) at a concentration of 1μg/cm^2^ for 12 hours (overnight) in complete media. Next day, all wells were serum starved for 3 hours. Cells were then maintained in media with 2% FBS or 2% media supplemented with EGF (25ng/ml). Cells were imaged for 48hrs on a time-interval of 15mins on Keyence BZ-X810. All pictures were stitched together to produce a video with speed of 14fps. For dynamic tracking, 5 single cells were analyzed on Keyence BZ-X810 software to generate Chemotaxis plot and to calculate movement and speed of cells.

### RNAi Sequences

All RNAi (siRNAs) used for study were ordered from Thermo Fisher Scientific Biosciences Inc. Lafayette, CO 80026, USA and transfections were done using Dharmafect1 as per the supplier’s instructions.

#### 1. Non-Targeting Control

UAAGGCUAUGAAGAGAUACAA

#### 2. UBQLN1

siU1^1^: GAAGAAAUCUCUAAACGUUUUUU

siU1^2^: GUACUACUGCGCCAAAUUUUU

### Statistical Analysis

All statistics were performed using GraphPad Prism 8 software. Unless otherwise specified, significance was determined by one-way ANOVA, using a cut off of p<0.05.

## Results

### Ubiquilin1 and Ubiquilin2 interact with EGFR

Our previous data described that UBQLN1 interacts with IGF1R. In this study, we aimed to explore the possibility that UBQLN proteins might interact with and regulate additional RTK family members, like EGFR. We first performed co-immunoprecipitation (IP) of UBQLNs to identify interacting proteins (Figure 1). We transiently transfected HEK293T cells with UBQLN1-FLAG or UBQLN2-FLAG constructs, then pulled down UBQLN-interacting proteins using anti-FLAG conjugated agarose beads. Western blot analysis of these FLAG-IP samples revealed that UBQLN1 and UBQLN2 interacted with total EGFR (Figure 1B). Like UBQLN1, UBQLN2 has an N-terminus UBL, a C-terminus UBA and four STI (STI1-4) domains, plus UBQLN2 has an additional, unique PXX (12 tandem repeats) domain (Figure 1A). To determine the interacting domain of UBQLN1, we created two constructs as described in Figure 1A and performed similar co-immunoprecipitation experiment. For one construct, we deleted the UBA domain of UBQLN1 (labeled 542X), and for the second construct we deleted the UBL domain (labeled 112X). Our results indicated that the UBL domain is the primary site of interaction with EGFR. Deletion of the UBA domain did not cause a loss of interaction between UBQLN1 and EGFR (Figure 1C).

### Ubiquilin1 regulates expression and activity of EGFR

Next, we investigated whether loss of UBQLN1 regulated EGFR expression and activity in lung adenocarcinoma cells (Figure 2). A549 cells were transiently transfected with non-targeting siRNA (NT) and two UBQLN1-specific siRNAs (Figure 2A). Cells were then serum starved (SS) to synchronize the cell cycle and remove all confounding growth factors present in serum. Then cells were treated with cycloheximide (CH) to prevent synthesis of new EGFR protein. Upon loss of UBQLN1, there was a decrease in total EGFR expression compared to the NT control. This decrease in total EGFR expression was significantly enhanced when the receptor was stimulated with EGF ligand. Next, we tested regulation of total EGFR when stimulated with different doses of EGF ligand (Figure 2B). There was significantly increased degradation of total EGFR in cells lacking UBQLN1 compared to controls at 10ng/ml, which was enhanced at 100ng/ml. Interestingly, expression of phosphorylated EGFR was not as sensitive to the loss of UBQLN1. Therefore, in cells lacking UBQLN1, there was an EGF dose-dependent degradation of EGFR.

**Figure 2:**
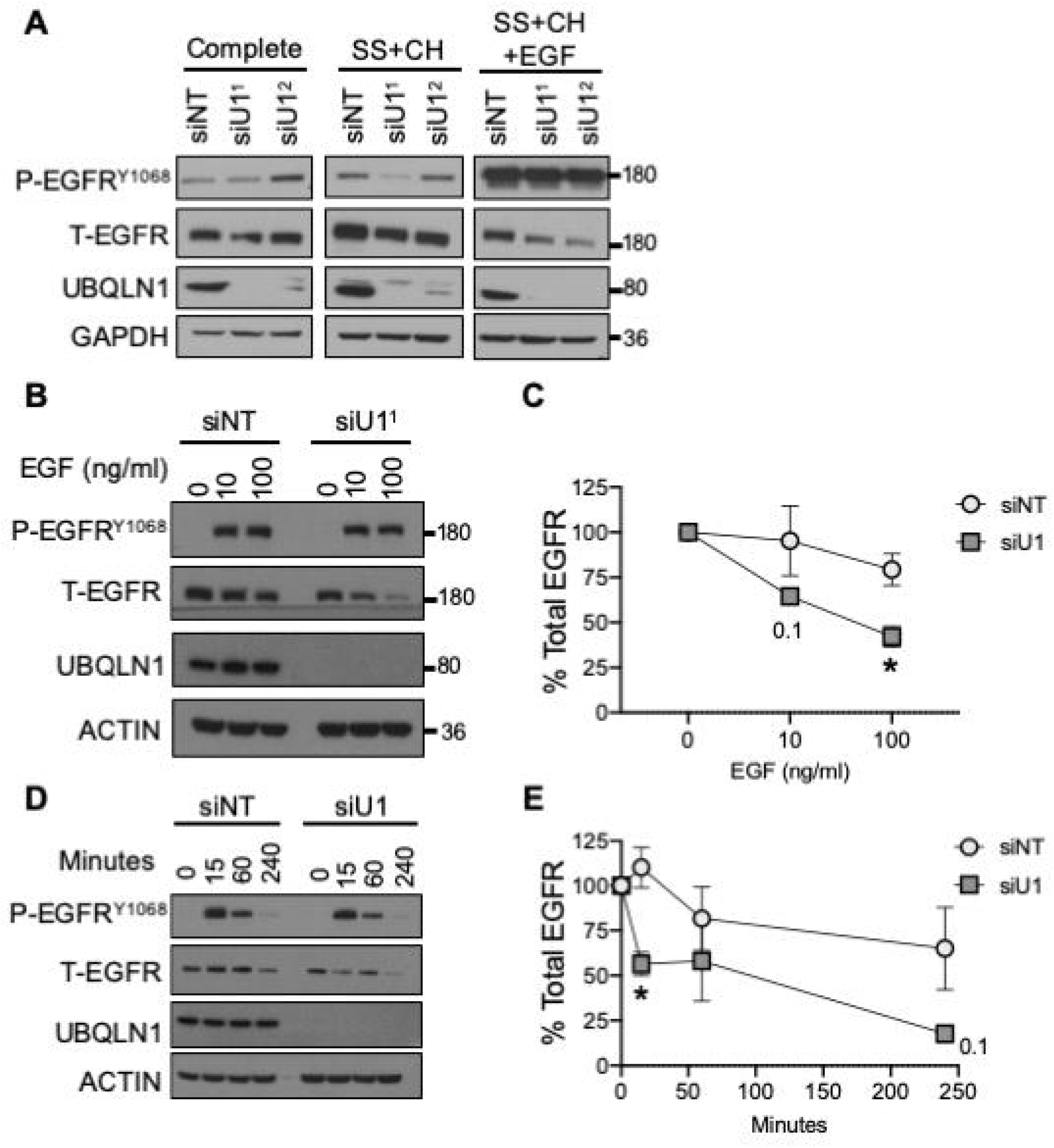
Ubiquilin1 regulates expression and activity of EGFR. **(A)** A549 cells were transiently transfected with two different siRNA’s for Ubiquilin1 (siU1^1^ and siU1^2^) along with non-targeting control (siNT). Cells were serum starved (SS) for 3 hours, incubated with a protein synthesis inhibitor, Cycloheximide for 1 hour, supplemented with EGF (50ng/ml) for 3 hours and analyzed by Western Blot. When stimulated with EGF, cells with loss of Ubiquilin1 demonstrated loss of total EGFR compared to controls. **(B)** A549 cells were transiently transfected with siRNA for Ubiquilin1, SS for 3 hours and stimulated with different does of EGF (10 and 100ng/ml) for 1 hour. Cells lacking Ubiquilin1 showed a dose-dependent loss of total EGFR, quantified in **(C)** n=3, Two-way ANOVA, p<0.05**. (D)** A549 cells were transiently transfected with siRNA for Ubiquilin1, SS for 3 hours and stimulated with EGF (50ng/ml) for indicated time points (0-240 minutes). As time passed, cells lacking Ubiquilin1 showed significantly increased loss of total EGFR compared to controls, quantified in **(E)** n=3, Two-way ANOVA, p<0.05.

Next, we performed experiments to determine temporal regulation of EGFR by UBQLN1 (Figure 2D, E). Lung adenocarcinoma cells were transfected with non-targeting or UBQLN1-specific siRNAs. Two days post-transfection, cells were serum starved for 3 hours, then stimulated with EGF (50ng/ml) for indicated time points. Upon stimulation with EGF, we observed degradation of total EGFR as time progressed. However, in cells lacking UBQLN1, there was significantly increased degradation compared to control as time progressed.

### Ubiquilin1-deficient cells exhibit increased cell viability and migration potential

EGFR proteins play a role in maintaining cell viability and stimulating migratory potential of lung adenocarcinoma cells. We investigated the influence of loss of UBQLN1 on these EGFR mediated processes (Figure 3). A549 cells were transiently transfected with siRNA for Ubiquilin1 and non-targeting control and cultured in different conditions as indicated (complete media, serum starvation (SS) for 3 hours, SS + EGF (50ng/ml), SS+EGF+Erlotinib, a phospho-EGFR inhibitor, 1uM). Cells were harvested after 3 hours and analyzed by Western Blot. When stimulated with EGF, Ubiquilin1 deficient cells showed almost complete loss of total EGFR and increased expression of phosphorylated EGFR, which was blocked by Erlotinib (Figure 3A). Next, A549 cells were transfected with non-targeting or UBQLN1-specific siRNAs, and cell viability was measured for four consecutive days using alamarBlue^™^ (Figure 3B). Consistent with our previous findings, loss of UBQLN1 resulted in increased cell growth in lung adenocarcinoma cell lines. Interestingly, there was an increase in the relative number of cells in UBQLN1 deficient cells stimulated with EGF, compared to controls. Next, we determined the effects of loss of UBQLN1 on cell migration (Figure 3C, D). Using a Transwell^™^ migration plate, we seeded A549 cells that had been transfected with either non-targeting or UBQLN1-specific siRNAs and cultured the cells under three conditions - serum-free media (SF); SF media supplemented with EGF; and SF media supplemented with EGF and Erlotinib, a phospho-EGFR inhibitor. Consistent with our previous data using this migration model, UBQLN1 deficient cells exhibited an approximately 3-fold increase in migration when stimulated with EGF. Erlotinib decreased migration of cells in both control and UBQLN1 deficient cells.

**Figure 3:**
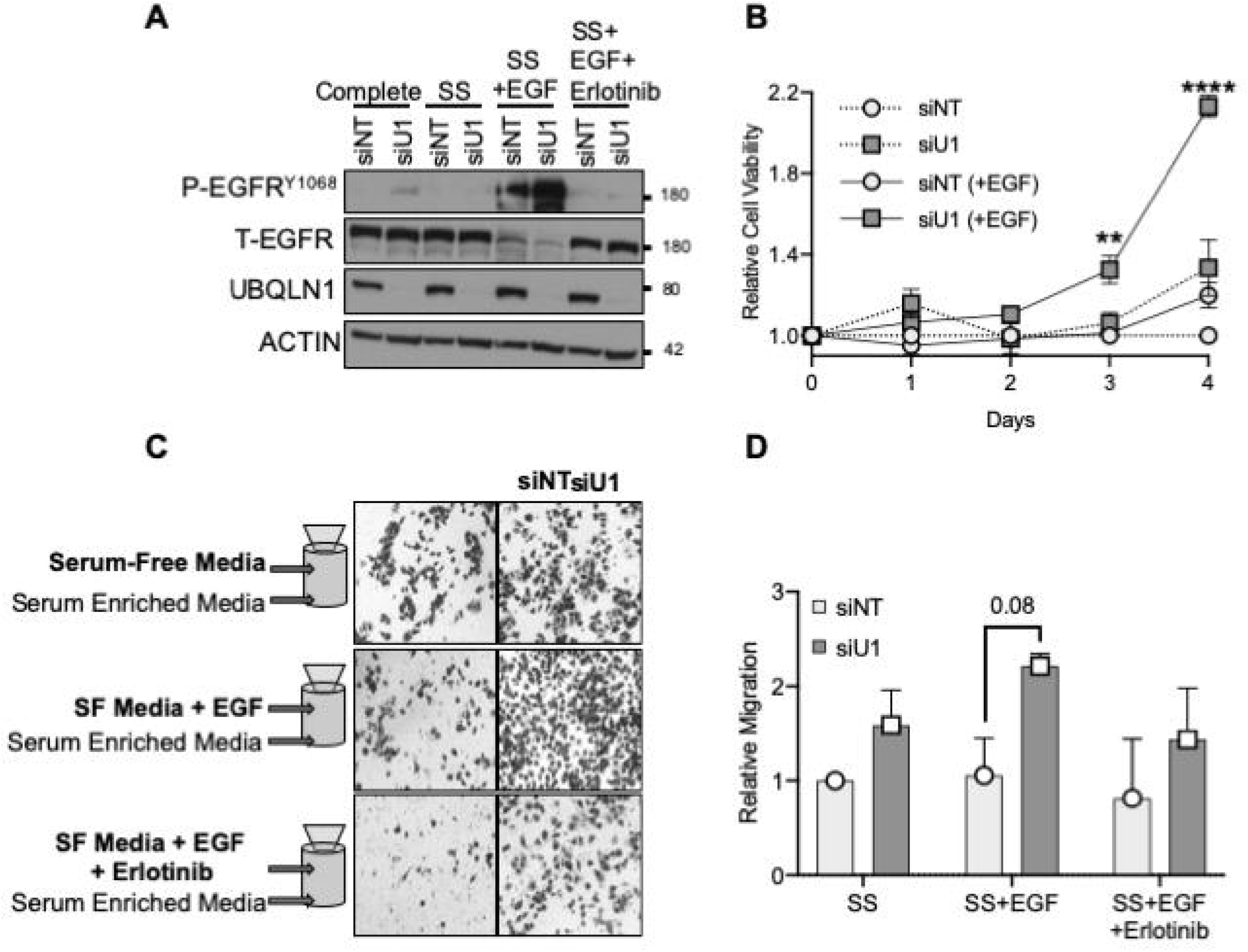
UBQLN1 deficient cells show increased cell viability and migration potential. **(A)** A549 cells were transiently transfected with siRNA for Ubiquilin1 and non-targeting control, and cultured in different conditions as indicated (complete media, serum starvation (SS) for 3 hours, SS + EGF (50ng/ml), SS+EGF+Erlotinib 1uM). Cells were harvested after 3 hours and analyzed by Western Blot. When stimulated with EGF, Ubiquilin1 deficient cells showed almost complete loss of total EGFR and increased phosphorylated EGFR. **(B)** A549 cells were transiently transfected with siRNA for Ubiquilin1 and non-targeting control. 12hrs post-transfection, cells reseeded in a 96-well plate for 12hrs (overnight). Cells were then serum starved for 3 hours followed by stimulation with EGF in 2% FBS and were cultured for 4 days. Alamar Blue readings were recorded every 24 hours and relative cell viability of UBQLN1 deficient cells were compared to control cells on each day. UBQLN1 deficient cells supplemented with EGF showed significantly increased viability compared to controls. One-way ANOVA, **p<0.01, ****p<0.-0001 **(C)** A549 cells were transiently transfected with siRNA for Ubiquilin1, seeded in a transwell setup to assess cell migration in response to EGF stimulation. Cells were cultured in the top chamber in one of 3 conditions – serum-free media, serum-free media supplemented with EGF and serum-free media supplemented with EGF and Erlotinib. Media supplemented with 10% FBS was used as chemo-attractant in the bottom chamber. At the end of 24 hours, cells were fixed and probed with HEMA 3 stain and data are quantified in **(D)**. Under all 3 conditions, UBQLN1 deficient cells demonstrated increased invasive behavior compared to controls. n=2, Two-way ANOVA, p<0.05.

### Loss of Ubiquilin1 results in increased cell movement and speed

We examined individual cell movement and speed to explore increased migratory potential in cells lacking UBQLN1 (Figure 4). We utilized live cell imaging equipped with image analysis software (Keyence). A549 lung adenocarcinoma cells were transfected with non-targeting or UBQLN1-specific siRNAs for 24 hours after which they were serum starved for another 24 hours. At this point, 10,000 cells per well (12-well plate) were seeded on a thin layer of commercial extracellular matrix (ECM) in the presence or absence of EGF (25ng/ml) to yield the following conditions: siNT (+/-) EGF and siUBQLN1 (+/-) EGF. Live cell images were captured for 48 hours using a Keyence microscope. All captured images were stitched together to make a representative video (Figure S1). As evident from the videos, loss of UBQLN1 resulted in increased movement of A549 cells which was further enhanced by EGF stimulation. These data are represented as chemotaxis plots (Figure 4A). Additionally, we quantified the distance traveled and the rate of travel (speed) for each cell (Figure 4B). These data show that cells lacking UBQLN1 travel further and move faster as compared to the non-targeting controls. These differences were enhanced in the presence of EGF.

**Figure 4:**
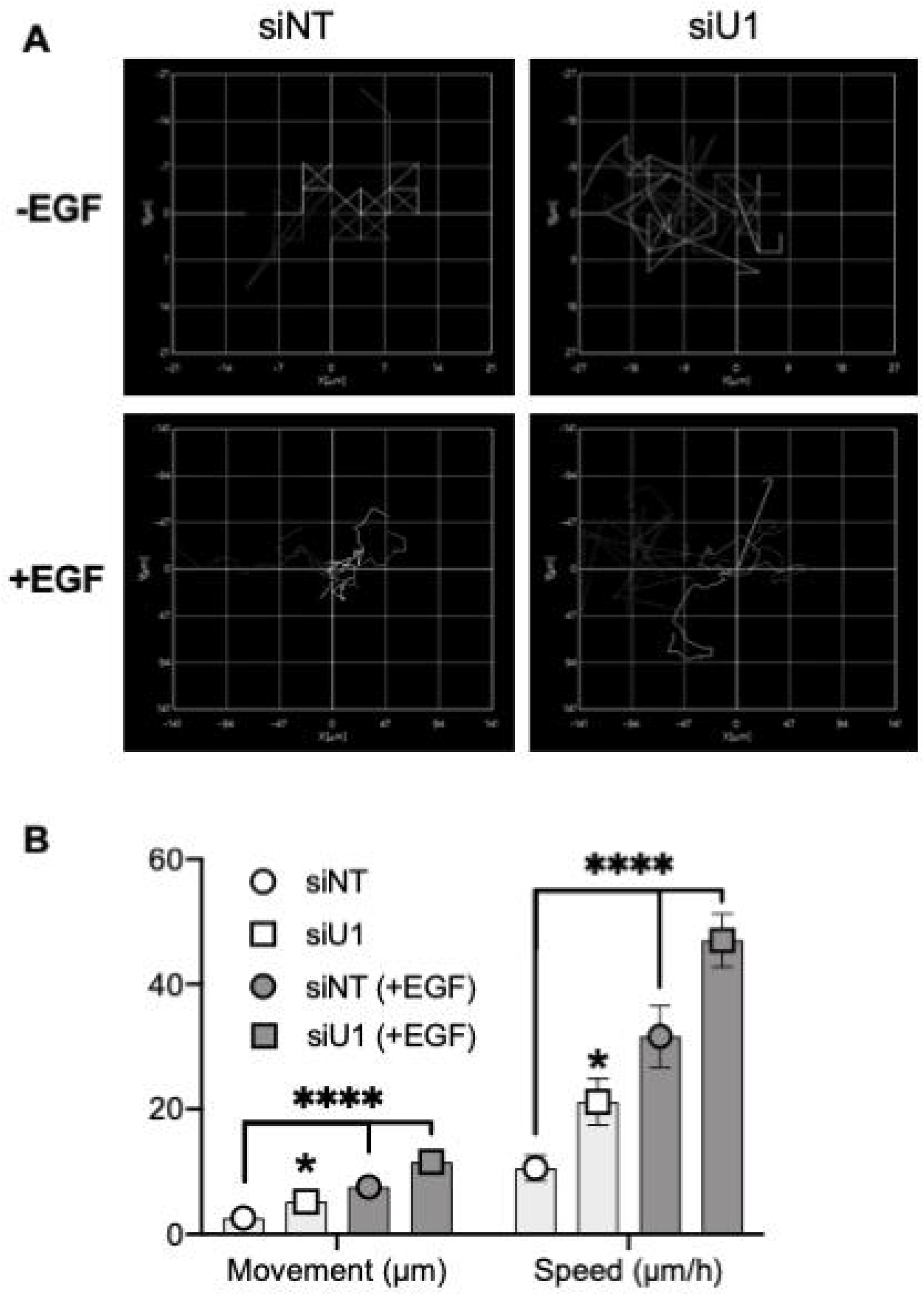
Loss of UBQLN1 results in increased cell movement and speed. **(A)** A549 cells were transiently transfected with siRNA for Ubiquilin1 and non-targeting control. 24hrs post-transfection, cells were reseeded in a plate coated with ECM (1μg/cm^2^) for 12hrs (overnight) in complete media. Cells were then serum starved for 3 hours followed by stimulation with EGF in 2% FBS. Cells were then imaged for 48hrs on a time-interval of 15mins. All pictures were stitched together to produce a video with speed of 14fps (Supplementary figure 1). **(B)** 5 single cells were analyzed on Keyence BZ-X810 to generate Chemotaxis plot. **(C)** Dynamic tracking were also used to calculate speed and movement of 5 single cells. One-way ANOVA, *p<0.05, ****p<0.0001.

## Discussion

UBQLN1 was reported to be lost in approximately fifty percent of non-small cell lung adenocarcinomas. Our group is interested in understanding how loss of function of UBQLN proteins contributes to the metastatic progression of human lung adenocarcinoma.(Z. Kurlawala, Dunaway, et al., 2017; Shah et al., 2015; Yadav et al., 2017). Previously we have shown that interaction between UBQLN1 with IGF1R results in stabilization of this receptor. When UBQLN1 is lost, it leads to increased phosphorylation of the auto-phosphorylation site on the IGF receptor, while total IGF1R levels decreased. Similarly, in this manuscript, we report that loss of UBQLN1 does not alter phosphorylated EGFR expression while causing a robust decrease in total EGFR expression. Additionally, we have reported dose and time dependent decreases in total IGF1R when stimulated with IGF ligand (Z. Kurlawala, Dunaway, et al., 2017). Likewise, in this study, lung adenocarcinoma cells lacking UBQLN1 also exhibit an enhanced degradation of EGFR when stimulated with EGF that is both dose and time dependent. These data suggest that loss of UBQLN1 accelerates both EGFR and IGF1R turnover in cancer cells. UBQLN1 might be crucial to maintain the stability of these RTKs (Z Kurlawala & Beverly, 2017), thus influencing pro-growth and pro-survival signaling pathways in lung adenocarcinoma cells.

We have previously demonstrated that downregulation of UBQLN1 leads to significantly increased expression of mesenchymal markers like Vimentin, Snail and ZEB1 indicating that UBQLN1 may play a role in suppression of metastasis in lung cancer (Shah et al., 2015). Additionally, knockdown of UBQLN1 by siRNA or mir155-mediated downregulation of UBQLN1 in lung cancer cells promoted an EMT-like phenotype. Data presented here further supports the role of UBQLN1 in proliferation and migration of cancer cells and this was exacerbated in the presence of EGF stimulation. These results, when considered in conjunction with our previous work with IGF1R, suggest that cells lacking UBQLN1, then stimulated with IGF or EGF, enhanced the metastatic potential of cancer cells(Z. Kurlawala, Dunaway, et al., 2017). Our results point to a destabilization of these RTKs when UBQLN1 is lost which further leads to an invasive phenotype in lung adenocarcinoma cells. In a related study, UBQLN4, another member of the UBQLN family, was shown to interact with RNF11, an E3 ubiquitin ligase of p21. Overexpression of UBQLN4 induced cellular senescence and cell cycle arrest in gastric cancer cells (S. Huang et al., 2019). These findings further support significance of UBQLN proteins in cancer progression and tumorigenesis. As reported here, cells that lack UBQLN1 show increased cell movement and speed captured with live cell imaging over multiple days. The overall movement and speed of these lung adenocarcinoma cells was further accelerated in the presence of EGF stimulation. RTK family of kinases play critical roles in progression of human lung cancer. In non-small cell lung cancer (NSCLC), IGFR and EGFR are overexpressed and UBQLNs are underexpressed and synergistically contribute to tumor development and progression (Guo et al., 2017; Oliveira, Schiffelers, Storm, Henegouwen, & Roovers, 2009). Collectively, these data indicate a critical role for UBQLNs in the normal proteolytic degradation of RTKs and loss of UBQLN function in cancer cells leads to aberrant RTK-mediated signaling, further enhancing the metastatic potential of these cancers. Downstream pathways activated by these cell surface receptors crosstalk and upon mutual activation lead to acquired resistance against EGFR-targeted drugs. We propose that targeting both EGF and IGF receptors at once might enhance anti-tumor efficacy and would be a promising approach for NSCLC therapies(Guo et al., 2017; Oliveira et al., 2009; Yeo et al., 2015). Being able to target both of these pathways simultaneously via their interaction with UBQLNs is an exciting avenue to explore.

Our lab is also interested in the phenomenon of inverse relation of cancer with neurodegenerative disorders. Single nucleotide polymorphisms (SNPs) in UBQLN1 gene are associated with late onset of Alzheimer’s Disease (AD) (Bertram et al., 2005). Additionally, mutations found in UBQLN2, provide a possible pathophysiological link for worse prognosis in amyotrophic lateral sclerosis/frontotemporal dementia (ALS/FTD)(Brettschneider et al., 2012; Renaud et al., 2019). After UBQLN1 was found to interact with the presenilin complex via a yeast-two hybrid screen, research efforts were quickly underway to establish the role that UBQLN family members might play in neurodegenerative diseases(Bertram et al., 2005; Brettschneider et al., 2012; Hiltunen et al., 2006; Mah et al., 2000; Renaud et al., 2019; Viswanathan et al., 2011). Perhaps, UBQLN family members inversely regulate outcomes in cancer and neurodegenerative diseases such that loss of function of UBQLN proteins in epithelial cells leads to a cancerous phenotype while in post-mitotic neurons leads to a degenerative phenotype. UBQLNs could play a regulatory role in presenilin complex assembly by directly, or indirectly, altering the stability of individual proteins found in these complexes thereby destabilizing the complex as a whole and leading to the pathogenesis of the disease state. Further experiments would be needed in order to establish the exact role.

Interestingly, some members of the BCL2 family, IGF1R, and EGFR are overexpressed in cancer but under-expressed in brains of AD patients (Shafi, 2016). Interestingly, each of these proteins, either in this study or in our previous work, have been shown to be stabilized by UBQLN1 (Beverly et al., 2012; Z Kurlawala & Beverly, 2017). In AD, β-amyloid aggregations accumulate in the brain. Studies have shown that inhibiting IGF1R and EGFR leads to significant reduction in β-amyloid aggregations (Wang et al. 2012; Gontier et al., 2015). Polyubiquitination of EGFR regulates its cellular location and stability, and this ubiquitination can be decreased by mutating the lysine residues without having an effect on the tyrosine kinase activity of EGFR(F. Huang, Kirkpatrick, Jiang, Gygi, & Sorkin, 2006). Thus, inhibition of RTKs could not only benefit certain subsets of cancer, it could also be beneficial in several neurodegenerative diseases(Gontier, George, Chaker, Holzenberger, & Aïd, 2015; Tavassoly et al., 2020; L. Wang et al., 2012). These studies further support our belief that UBQLNs might be regulating multiple diseases indirectly.

In conclusion, the UBA-UBL domain-containing family of UBQLNs facilitate normal proteasomal degradation, and substrate selection by UBQLNs is critical in cancer and neurodegenerative diseases. Our IP data provides evidence for interaction of both UBQLN1 and UBQLN2 (shares more than 75% homology with UBQLN1) with EGFR. Taken together with our previous findings, both IGFRs and EGFR were detected as interacting partners of UBQLN1. The lung adenocarcinoma cells lacking UBQLN1 exhibit enhanced degradation of both EGFR and IGF1R, especially in the presence of their respective ligands. Thus, UBQLN1 is important in maintaining stability of RTKs, like EGFR and IGF1R. Loss of UBQLN1 alters the normal degradative fate of these receptors leading to downstream alterations in signaling that result in a cellular phenotype with enhanced proliferation, migration and movement of the cells. The RTK family of proteins are highly regulated in both cancer and neurodegenerative diseases and this sets the stage for the development of directed therapies. Our work with UBQLNs indicates that they are uniquely poised as therapeutic targets given their role in regulating the stability of multiple members of the RTK family, and this regulation might be inversely impacted in cancer versus neurodegenerative diseases.

## List of Abbreviations

AD: Alzheimer’s disease
UBQLNs: Ubiquilin family of adapter proteins
UBA: Ubiquitin-associated domain
UBL: Ubiquitin-like domain
APP: β-amyloid precursor protein
ER: Endoplasmic reticulum
EMT: Epithelial to mesenchymal transition
BCL2: B-cell lymphoma 2
RTK: Receptor tyrosine kinase
IGFR: Insulin-like growth factor receptor
EGFR: Epidermal growth factor receptor
ALS: Amyotrophic lateral sclerosis
NSCLC: Non-small cell lung cancer
EGF: Epidermal growth factor (ligand).

## Declarations

### Consent for publication

Not applicable

### Availability of data and materials

The datasets used and/or analyzed during the current study are available from the corresponding author upon reasonable request.

### Authors’ contributions

KS, ZK, RD and PPS did the experiments. KS, ZK, LJS and LJB conceived studies, did the analysis and wrote the manuscript. All authors have read and approved the manuscript.

## Acknowledgements

We are grateful to the Beverly and Siskind lab members, especially Lavona Casson and Mark Doll, for continuous support and sharing of resources and insight.

## References

Bertram, L., Hiltunen, M., Parkinson, M., Ingelsson, M., Lange, C., Ramasamy, K.,… Tanzi, R. E. (2005). Family-based association between Alzheimer’s disease and variants in UBQLN1. N Engl J Med, 352(9), 884–894. doi:10.1056/NEJMoa042765

Beverly, L. J., Lockwood, W. W., Shah, P. P., Erdjument-Bromage, H., & Varmus, H. (2012). Ubiquitination, localization, and stability of an anti-apoptotic BCL2-like protein, BCL2L10/BCLb, are regulated by Ubiquilin1. Proc Natl Acad Sci U S A, 109(3), E119–126. doi:10.1073/pnas.1119167109

Brettschneider, J., Van Deerlin, V. M., Robinson, J. L., Kwong, L., Lee, E. B., Ali, Y. O.,… Elman, L. (2012). Pattern of ubiquilin pathology in ALS and FTLD indicates presence of C9ORF72 hexanucleotide expansion. Acta Neuropathol, 123(6), 825–839.

Feng, P., Scott, C. W., Cho, N. H., Nakamura, H., Chung, Y. H., Monteiro, M. J., & Jung, J. U. (2004). Kaposi’s sarcoma-associated herpesvirus K7 protein targets a ubiquitin-like/ubiquitin-associated domain-containing protein to promote protein degradation. Mol Cell Biol, 24(9), 3938–3948. Retrieved from https://www.ncbi.nlm.nih.gov/pubmed/15082787

Gontier, G., George, C., Chaker, Z., Holzenberger, M., & Aïd, S. (2015). Blocking IGF signaling in adult neurons alleviates Alzheimer’s disease pathology through amyloid-β clearance. Journal of Neuroscience, 35(33), 11500–11513.

Guo, X.-F., Zhu, X.-F., Cao, H.-Y., Zhong, G.-S., Li, L., Deng, B.-G.,… Zhen, Y.-S. (2017). A bispecific enediyne-energized fusion protein targeting both epidermal growth factor receptor and insulin-like growth factor 1 receptor showing enhanced antitumor efficacy against non-small cell lung cancer. Oncotarget, 8(16), 27286.

Hiltunen, M., Lu, A., Thomas, A. V., Romano, D. M., Kim, M., Jones, P. B.,… Berezovska, O. (2006). Ubiquilin 1 modulates amyloid precursor protein trafficking and Aβ secretion. Journal of Biological Chemistry, 281(43), 32240–32253.

Huang, F., Kirkpatrick, D., Jiang, X., Gygi, S., & Sorkin, A. (2006). Differential regulation of EGF receptor internalization and degradation by multiubiquitination within the kinase domain. Molecular Cell, 21(6), 737–748.

Huang, S., Li, Y., Yuan, X., Zhao, M., Wang, J., Li, Y.,… Huang, C. (2019). The UbL-UBA Ubiquilin4 protein functions as a tumor suppressor in gastric cancer by p53-dependent and p53-independent regulation of p21. Cell Death Differ, 26(3), 516–530. doi:10.1038/s41418-018-0141-4

Kleijnen, M. F., Alarcón, R. M., & Howley, P. M. (2003). The ubiquitin-associated domain of hPLIC-2 interacts with the proteasome. Molecular biology of the cell, 14(9), 3868–3875.

Kleijnen, M. F., Shih, A. H., Zhou, P., Kumar, S., Soccio, R. E., Kedersha, N. L.,… Howley, P. M. (2000). The hPLIC proteins may provide a link between the ubiquitination machinery and the proteasome. Molecular Cell, 6(2), 409–419.

Kurlawala, Z., & Beverly, L. (2017). Ubiquilin Proteins are critical adaptors that regulate proteostasis. Journal of Cell Signaling, 2, 145.

Kurlawala, Z., Dunaway, R., Shah, P. P., Gosney, J. A., Siskind, L. J., Ceresa, B. P., & Beverly, L. J. (2017). Regulation of insulin-like growth factor receptors by Ubiquilin1. Biochem J, 474(24), 4105–4118. doi:10.1042/BCJ20170620

Kurlawala, Z., Shah, P. P., Shah, C., & Beverly, L. J. (2017). The STI and UBA Domains of UBQLN1 are Critical Determinants of Substrate Interaction and Proteostasis. J Cell Biochem. doi:10.1002/jcb.25880

Lee, D. Y., Arnott, D., & Brown, E. J. (2013). Ubiquilin4 is an adaptor protein that recruits Ubiquilin1 to the autophagy machinery. EMBO Rep, 14(4), 373–381. doi:10.1038/embor.2013.22

Lim, P. J., Danner, R., Liang, J., Doong, H., Harman, C., Srinivasan, D.,… Monteiro, M. J. (2009). Ubiquilin and p97/VCP bind erasin, forming a complex involved in ERAD. J Cell Biol, 187(2), 201–217. doi:10.1083/jcb.200903024

Lurje, G., & Lenz, H.-J. (2009). EGFR signaling and drug discovery. Oncology, 77(6), 400–410.

Mah, A. L., Perry, G., Smith, M. A., & Monteiro, M. J. (2000). Identification of ubiquilin, a novel presenilin interactor that increases presenilin protein accumulation. J Cell Biol, 151(4), 847–862. Retrieved from https://www.ncbi.nlm.nih.gov/pubmed/11076969

Marin, I. (2014). The ubiquilin gene family: evolutionary patterns and functional insights. BMC Evol Biol, 14, 63. doi:10.1186/1471-2148-14-63

Massey, L. K., Mah, A. L., Ford, D. L., Miller, J., Liang, J., Doong, H., & Monteiro, M. J. (2004). Overexpression of ubiquilin decreases ubiquitination and degradation of presenilin proteins. J Alzheimers Dis, 6(1), 79–92. Retrieved from http://www.ncbi.nlm.nih.gov/pubmed/15004330

N’Diaye, E. N., Hanyaloglu, A. C., Kajihara, K. K., Puthenveedu, M. A., Wu, P., von Zastrow, M., & Brown, E. J. (2008). The ubiquitin-like protein PLIC-2 is a negative regulator of G protein-coupled receptor endocytosis. Mol Biol Cell, 19(3), 1252–1260. doi:10.1091/mbc.E07-08-0775

N’Diaye, E. N., Kajihara, K. K., Hsieh, I., Morisaki, H., Debnath, J., & Brown, E. J. (2009). PLIC proteins or ubiquilins regulate autophagy-dependent cell survival during nutrient starvation. EMBO reports, 10(2), 173–179.

Oliveira, S., Schiffelers, R., Storm, G., Henegouwen, P., & Roovers, R. (2009). Crosstalk between epidermal growth factor receptor-and insulin-like growth factor-1 receptor signaling: implications for cancer therapy. Current cancer drug targets, 9(6), 748–760.

Porta, R., Sanchez-Torres, J., Paz-Ares, L., Massuti, B., Reguart, N., Mayo, C.,… Salinas, P. (2011). Brain metastases from lung cancer responding to erlotinib: the importance of EGFR mutation. European Respiratory Journal, 37(3), 624–631.

Renaud, L., Picher-Martel, V., Codron, P., & Julien, J. P. (2019). Key role of UBQLN2 in pathogenesis of amyotrophic lateral sclerosis and frontotemporal dementia. Acta Neuropathol Commun, 7(1), 103. doi:10.1186/s40478-019-0758-7

Saliba, R. S., Pangalos, M., & Moss, S. J. (2008). The ubiquitin-like protein Plic-1 enhances the membrane insertion of GABAA receptors by increasing their stability within the endoplasmic reticulum. J Biol Chem, 283(27), 18538–18544. doi:10.1074/jbc.M802077200

Shafi, O. (2016). Inverse relationship between Alzheimer’s disease and cancer, and other factors contributing to Alzheimer’s disease: a systematic review. BMC neurology, 16(1), 236.

Shah, P. P., Lockwood, W. W., Saurabh, K., Kurlawala, Z., Shannon, S. P., Waigel, S.,… Beverly, L. J. (2015). Ubiquilin1 represses migration and epithelial-to-mesenchymal transition of human non-small cell lung cancer cells. Oncogene, 34(13), 1709–1717. doi:10.1038/onc.2014.97

Sun, Q., Liu, T., Yuan, Y., Guo, Z., Xie, G., Du, S.,… Chen, L. (2015). MiR-200c inhibits autophagy and enhances radiosensitivity in breast cancer cells by targeting UBQLN1. Int J Cancer, 136(5), 1003–1012. doi:10.1002/ijc.29065

Tavassoly, O., Sato, T., & Tavassoly, I. (2020). Inhibition of Brain EGFR Activation: A Novel Target in Neurodegenerative Diseases and Brain Injuries. Molecular Pharmacology.

Viswanathan, J., Haapasalo, A., Bottcher, C., Miettinen, R., Kurkinen, K. M., Lu, A.,… Hiltunen, M. (2011). Alzheimer’s disease-associated ubiquilin-1 regulates presenilin-1 accumulation and aggresome formation. Traffic, 12(3), 330–348. doi:10.1111/j.1600-0854.2010.01149.x

Wang, L., Chiang, H.-C., Wu, W., Liang, B., Xie, Z., Yao, X.,… Zhong, Y. (2012). Epidermal growth factor receptor is a preferred target for treating Amyloid-β– induced memory loss. Proceedings of the National Academy of Sciences, 109(41), 16743–16748.

Wang, Y., Lu, J., Zhao, X., Feng, Y., Lv, S., Mu, Y.,… Li, Y. (2015). Prognostic significance of Ubiquilin1 expression in invasive breast cancer. Cancer Biomark, 15(5), 635–643. doi:10.3233/CBM-150503

Wu, A. L., Wang, J., Zheleznyak, A., & Brown, E. J. (1999). Ubiquitin-related proteins regulate interaction of vimentin intermediate filaments with the plasma membrane. Mol Cell, 4(4), 619–625. Retrieved from http://www.ncbi.nlm.nih.gov/pubmed/10549293

Yadav, S., Singh, N., Shah, P. P., Rowbotham, D. A., Malik, D., Srivastav, A.,… Beverly, L. J. (2017). MIR155 Regulation of Ubiquilin1 and Ubiquilin2: Implications in Cellular Protection and Tumorigenesis. Neoplasia, 19(4), 321–332. doi:10.1016/j.neo.2017.02.001

Yeo, C. D., Park, K. H., Park, C. K., Lee, S. H., Kim, S. J., Yoon, H. K.,… Kim, T.-J. (2015). Expression of insulin-like growth factor 1 receptor (IGF-1R) predicts poor responses to epidermal growth factor receptor (EGFR) tyrosine kinase inhibitors in non-small cell lung cancer patients harboring activating EGFR mutations. Lung cancer, 87(3), 311–317.

Zwick, E., Bange, J., & Ullrich, A. (2001). Receptor tyrosine kinase signalling as a target for cancer intervention strategies. Endocrine-related cancer, 8(3), 161–173.

